# Simultaneous in vitro simulation of multiple antimicrobial agents with different elimination half-lives in a pre-clinical infection model

**DOI:** 10.1101/2020.12.21.405464

**Authors:** Iordanis Kesisoglou, Brianna M Eales, Kimberly R. Ledesma, Paul R. Merlau, Vincent H Tam, Weiqun Wang, Michael Nikolaou

## Abstract

Combination therapy for treatment of multi-drug resistant bacterial infections is becoming increasingly common. In vitro testing of drug combinations under realistic pharmacokinetic conditions is needed before a corresponding combination is eventually put into clinical use. The current standard for in vitro simulation of the pharmacokinetics of two drugs with distinct elimination half-lives cannot be extended for combinations of three or more agents, posing a growing need. To address that need we develop a general method to design an in vitro model for simultaneous simulation of the kinetics of an arbitrary number of *N* drugs with different half-lives. The method developed entails two possible configurations: (a) An in-series configuration, which generalizes the standard two-drug design for *N* drugs and offers additional flexibility even for two drugs, and (b) an in-parallel configuration, which is new, and offers yet additional flexibility over the in-series configuration. Corresponding design equations for sizing and operation of each configuration are rigorously developed for immediate use by experimenters. These equations were used for experimental verification using a combination of three antibiotics with distinctly different half-lives (meropenem, ceftazidime, and levofloxacin). While experimental verification involved antibiotics, the method is applicable to any anti-infective or anti-cancer drugs with distinct elimination pharmacokinetics. With increasing importance of in vitro simulation of the kinetics of an arbitrary number of drugs in combination, the methods developed here are an important new tool for the design of such in vitro models.

## Introduction

Treatment of challenging bacterial infections by simultaneous use of two or more antimicrobial agents, known as combination therapy, has become common in recent years [1–3] as multi-drug resistant bacteria become increasingly prevalent [4–8]. Whether combination of certain agents has synergistic or antagonistic therapeutic effects must be realistically assessed before use, for the safety and successful treatment of patients [1,9,2]. In vitro testing of the effectiveness of agent combinations under realistic pharmacokinetic conditions is useful before a corresponding combination is eventually put into clinical use [3,10,11]. To simulate in vitro the pharmacokinetics of two agents with distinct elimination half-lives, Blaser [12] proposed a design method that has long been the standard for an experimental set-up which allows the concentration of each of the two agents in a common solution follows its own exponential decline over time, with a corresponding half-life. While immediately useful for a combination of two agents, Blaser’s results cannot be used for combinations of three or more agents, a situation that is becoming increasingly more common with multi-drug resistant bacteria [1–3]. Indeed, improved therapeutic effects have been observed from combination of ertapenem, meropenem, and colistin compared to double carbapenem alone [3], as well as from combining up to three of the following antibiotics: an aminoglycoside, an anti-pseudomonal beta-lactam, colistin, a fluoroquinolone, a macrolide, or rifampin [2]. Given the growing importance of combination therapy entailing three or more agents, this paper presents the development and experimental validation of a general method for designing in vitro simulations of the pharmacokinetics of an arbitrary number, *N*, of antimicrobial agents in combination, each agent with its own distinct elimination half-life. Formulas are developed for the design of an in vitro simulation system that extends Blaser’s configuration and offer substantially higher flexibility even for the case of two agents. In addition, a novel configuration with some advantageous features and corresponding design formulas are developed.

In the rest of the paper we first present relevant theoretical background summarizing experiment design for in vitro simulation of the pharmacokinetics of two agents. Subsequently, in Materials and Methods we present the theoretical and experimental framework within which we develop and test the proposed experiment design method for *N* ≥ 3 agents. The results of the proposed approach are presented next in the form of design equations. The design approach is illustrated experimentally on a hollow-fiber system [13–15]. A discussion of the obtained theoretical and experimental results follows. Finally, conclusions are drawn and recommendations for further development are made.

## Theoretical

### Design of experiments for in vitro simulation of distinct pharmacokinetics for two agents in combination

While in vivo bacterial response to antimicrobial agents can be assessed in animal infection models, there are uncertainty and risk when translating data from experiments on animals to human treatment, due to differences between animals and humans in agent pharmacokinetics [1,16,11]. To address this issue through in vitro simulation, a hollow-fiber (HF) system (Fig. 1) can be used to assess the response of bacterial populations exposed to antimicrobial agents using clinically relevant dosing regimens, commensurate with agent pharmacokinetics in humans [17,18]. Bacteria are exposed to nutrients and agents but are restrained from leaving the hollow fiber cartridge.

**Fig. 1.**
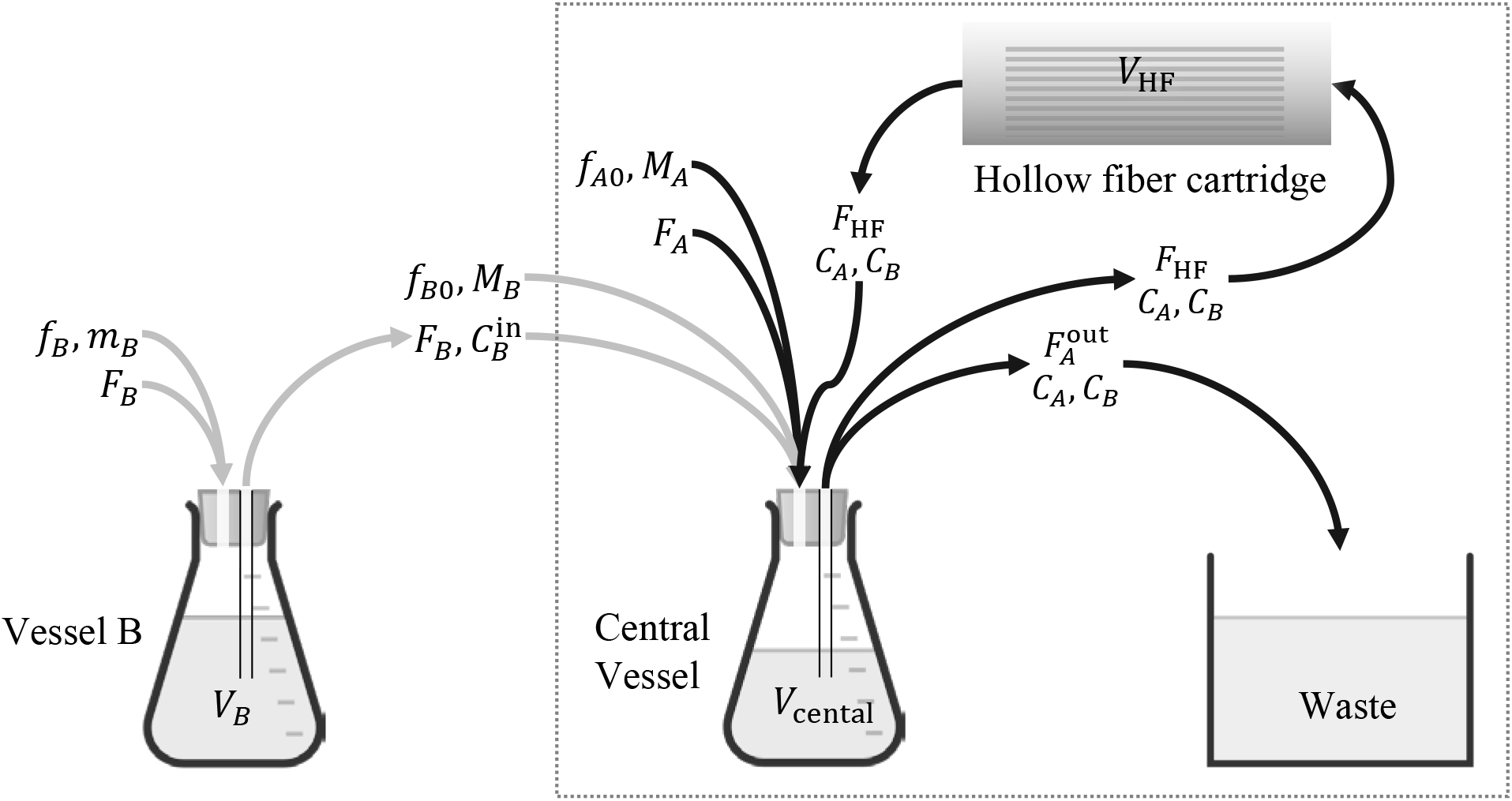
A hollow-fiber system for in vitro simulation of clinically relevant pharmacokinetics of antimicrobial agents. The system inside the dotted frame is suitable for a single agent, A; addition of vessel B in series with the central vessel renders it suitable for two agents, A, B. Volumetric flow rates F_A_, F_B_ entering vessels B and central are sterile growth media. Mass flow rates f_B_, f_B0_, and f_A0_ refer to bolus injections m_B_, M_B_, M_A_ of corresponding agents. With appropriate selection of 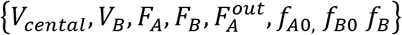 [12], the entire system is suitable for handling simultaneously two agents, A and B, each following its own elimination pharmacokinetics as they go through the hollow-fiber cartridge. For corresponding j, the notation is as follows: Fj = volumetric flow rate containing agent j = A, B. F_HF_ = internal circulation flow rate through the hollow fiber cartridge. 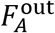 = volumetic flow rate from the central vessel to waste. f_j_ = volumetric flow rate of injection into vessel j containing agent j = B (Fig. 3). f_j0_ = volumetric flow rate of injection into the central vessel, j = A, B. V_j_ = liquid volume in vessel j = B. V_central_ = liquid volume in central vessel. V_HF_ = liquid volume in hollow fiber cartridge. C_j_ = concentration of agent j = A, B. 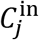 = concentration of agent j = B into the central vessel. mj = mass of agent j = B in bolus injection to vessel j = B. Mj = mass of agent j = A, B in bolus injection to central vessel.

Design of a hollow-fiber system for one or two agents, as shown in Fig. 1, refers to selecting the agent and broth flow rates as well as liquid volumes in each vessel, such that the profiles of *C_A_*(*t*) or {*C_A_*(*t*), *C_B_*(*t*)}, for one or two agents respectively, follow prescribed pharmacokinetics. Such pharmacokinetics typically corresponds to exponential decline of agent concentration over time after an initial peak, as captured by the following equation:

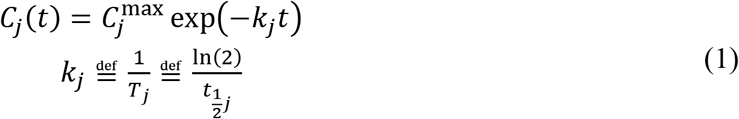

over time 0 ≤ *t < P*, where *k_j_* is the elimination rate constant of agent *j*; *T_j_* is the elimination time constant of agent *j*; 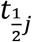 is the elimination half-life of agent *j*; *j* = *A* or *j* = *A, B*, for one or two agents, respectively. As explained by Blaser [17,12] regarding the configuration shown in Fig. 1, the exponentially declining concentration profiles in eqn. (1) result from impulse bolus injections into two vessels rather than into one, as a single vessel, with a single time constant equal to liquid volume over total flow rate, would be inadequate for creating distinct concentration profiles for two agents with different half-lives. Note that the period, *P*, between two successive injections is assumed in [12] to be large enough to ensure that *C_j_*(*P*) ≈ 0 at the time immediately before each new injection, as shown in Fig. 2.

**Fig. 2.**
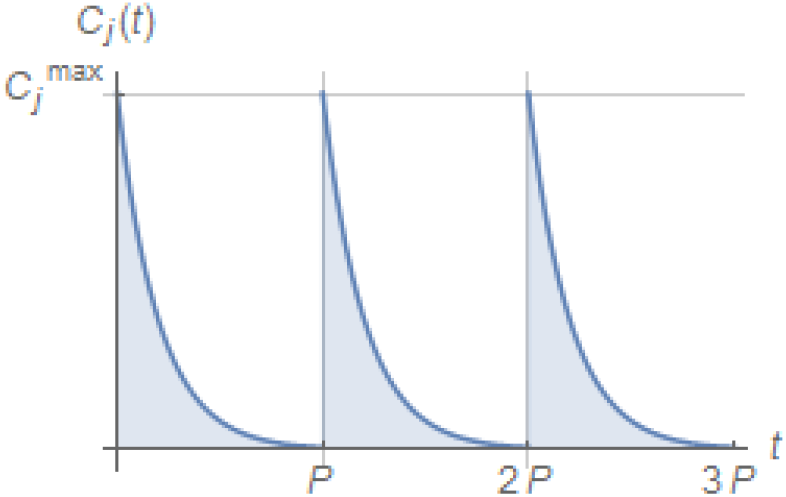
Pharmacokinetic profile corresponding to periodic injections of an antimicrobial agent at times nP, n = 0,1,2,…, with subsequent exponential decline of agent concentration 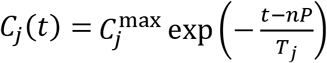 as shown in eqn. (1).

A summary of the design procedure for the hollow-fiber system of Fig. 1, accommodating either one or two agents, is shown in Table 1 (input data to the design) and Table 2 (output data of the design).

**Table 1.** Input data in the design procedure for the hollow-fiber system in Fig. 1 with one or two antimicrobial agents, following Blaser [17,12]

| Parameter description | Parameter name | Agent, $j$ |
| --- | --- | --- |
| Half-life of antimicrobial agent $j$ | $t_{\frac{1}{2}j}$ | $A$ or $\{A, B\}$ |
| Period between two successive injections | $P$ | $A$ or $\{A, B\}$ |
| Peak concentration of antimicrobial agent $j$ in the central vessel at injection time | $C_j^{\max}$ | $A$ or $\{A, B\}$ |
| Liquid volume in combined central vessel and hollow-fiber cartridge | $V_A \stackrel{\text{def}}{=} V_{\text{central}} + V_{\text{HF}}$ | |

**Table 2.** Summary of design procedure given data in Table 1 for the hollow-fiber system in Fig. 1 with one or two antimicrobial agents, following Blaser [17,12].

| Parameter description | Parameter name and design equation | Agent, $j$ |
| --- | --- | --- |
| Total volumetric flow rate out of central vessel, $F_A^{\text{out}}$ | $F_A^{\text{out}} = V_A k_A$ | |
| Volumetric flow rate of B into and out of vessel B, $F_B$ | $F_B, F_A, V_B$<br>are non-zero solutions of<br>$V_B = \frac{F_A^{\text{out}} - F_A}{F_A} V_A$ $F_B = k_B V_B$ $F_A^{\text{out}} = F_A + F_B$ | |
| Volumetric flow rate of A into central vessel, $F_A$ | | |
| Liquid volume in vessel B, $V_B$ | | |
| Mass, $M_j$, of impulse bolus injection into the central vessel, approximated by a pulse of short duration $\Delta t$ (Fig. 3) | $M_j = C_j^{\max} V_A$ | $A$ or $\{A, B\}$ |
| Mass, $m_B$, of impulse bolus injection into the vessel B, approximated by a pulse of short duration $\Delta t$ (Fig. 3) | $m_B = \frac{V_B}{V_A} M_B$ | |

Note that $f_{j0}(t)$ , $j = A, B$ , and $f_B(t)$ , $f_{B0}(t)$ in and Fig. 1 refer to corresponding mass flow rate approximation of impulses by pulses of short duration $\Delta t$ (Fig. 3) and of corresponding amplitudes, for appropriate bolus injections as follows:

Note that *f*_*j*0_(*t*) = *A, B*, and *f_B_*(*t*),*f*_*B*0_(*t*) in and Fig. 1 refer to corresponding mass flow rate approximation of impulses by pulses of short duration Δ*t* (Fig. 3) and of corresponding amplitudes, for appropriate bolus injections as follows:

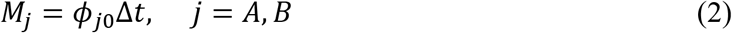

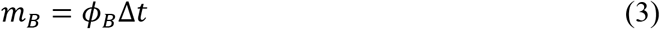

where *ϕ* denotes pulse amplitude, as shown in Fig. 3, with specific *ϕ_A0_, ϕ_B0_, ϕ_B_* referring to Fig. 1. These bolus injections may occur periodically at times *nP,n* = 0,1,2,…, in correspondence with Fig. 2.

**Fig. 3.**
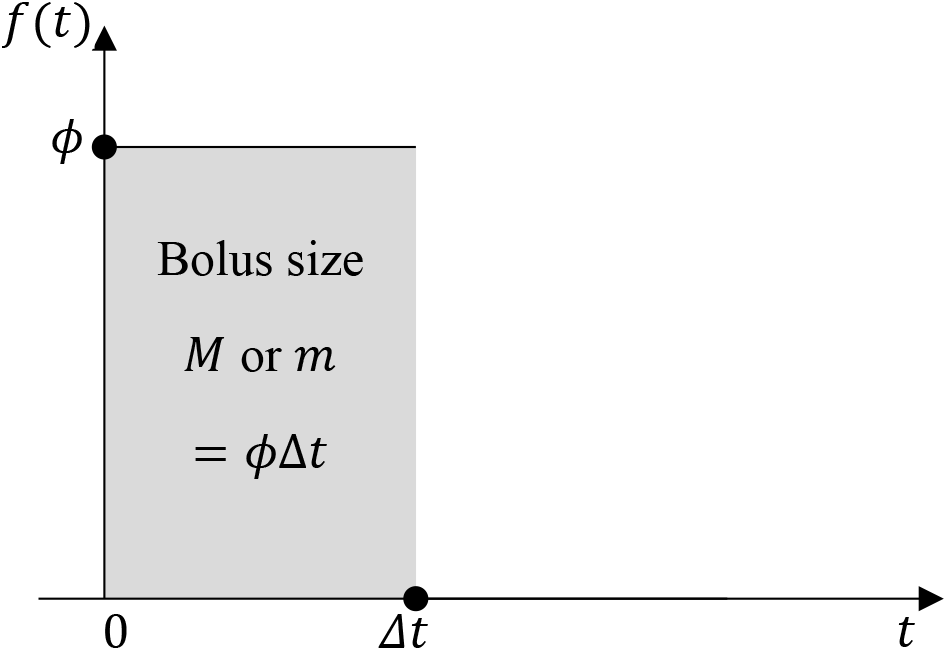
Mass flow rate f(t) approximating a bolus impulse of mass m or M by a pulse of high amplitude ϕ and short duration Δt.

Finally, note that in the case of two agents, *A, B*, the agent with the smaller half-life must be accommodated in the central vessel, i.e. the experimental set-up should make certain that

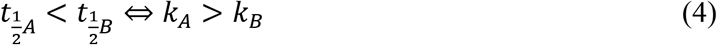

because the formulas in Table 2 immediately lead to

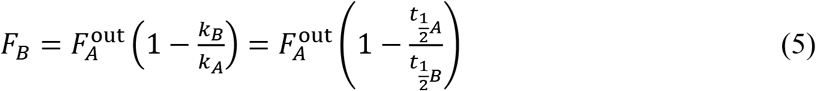

which must be positive, thus implying eqn. (4).

As already mentioned, the above method summarized in Table 1 and Table 2 is suitable for up to two antimicrobial agents. In the rest of the paper we present the development of a method suitable for an arbitrary number of agents, *N*, while also relaxing some of the restrictions placed in [12].

## Materials and Methods

### Set-up for in vitro simulation of distinct pharmacokinetics for *N* agents

As Blaser [12] argued, for the case of two agents, a vessel B in series with the central vessel must be included (Fig. 1) for an agent B to exhibit a concentration profile distinct from A as it goes through the hollow-fiber cartridge. Because Blaser’s design is limited to two agents, we develop here the following two new designs, suitable for any number of agents:

a. Starting with Fig. 1, which is suitable for two agents, Fig. 4 shows a configuration of a hollow-fiber system capable of handling three agents, A, B, C. In that configuration, a vessel for agent C is used *in series* with (i.e. feeding to) the central vessel, in the same fashion as vessel B. Four or more agents can be similarly handled by adding more vessels, all feeding to the central vessel. The central vessel in the configuration of Fig. 4 remains dedicated to handling agent A, through direct injection of bolus *M_A_* and broth inlet flow rate *F_A_*.
b. Departing from the in-series configuration, we propose the novel experiment configuration shown in Fig. 5. In that configuration, injections of A, B, C are used into three corresponding vessels *in parallel* to each other. The outlet streams of these vessels are fed to the central vessel. Separate injections of A, B, C take place into the central vessel as well. We will demonstrate in the Results section that this configuration has some interesting properties that distinguish it from the in-series design.

**Fig. 4.**
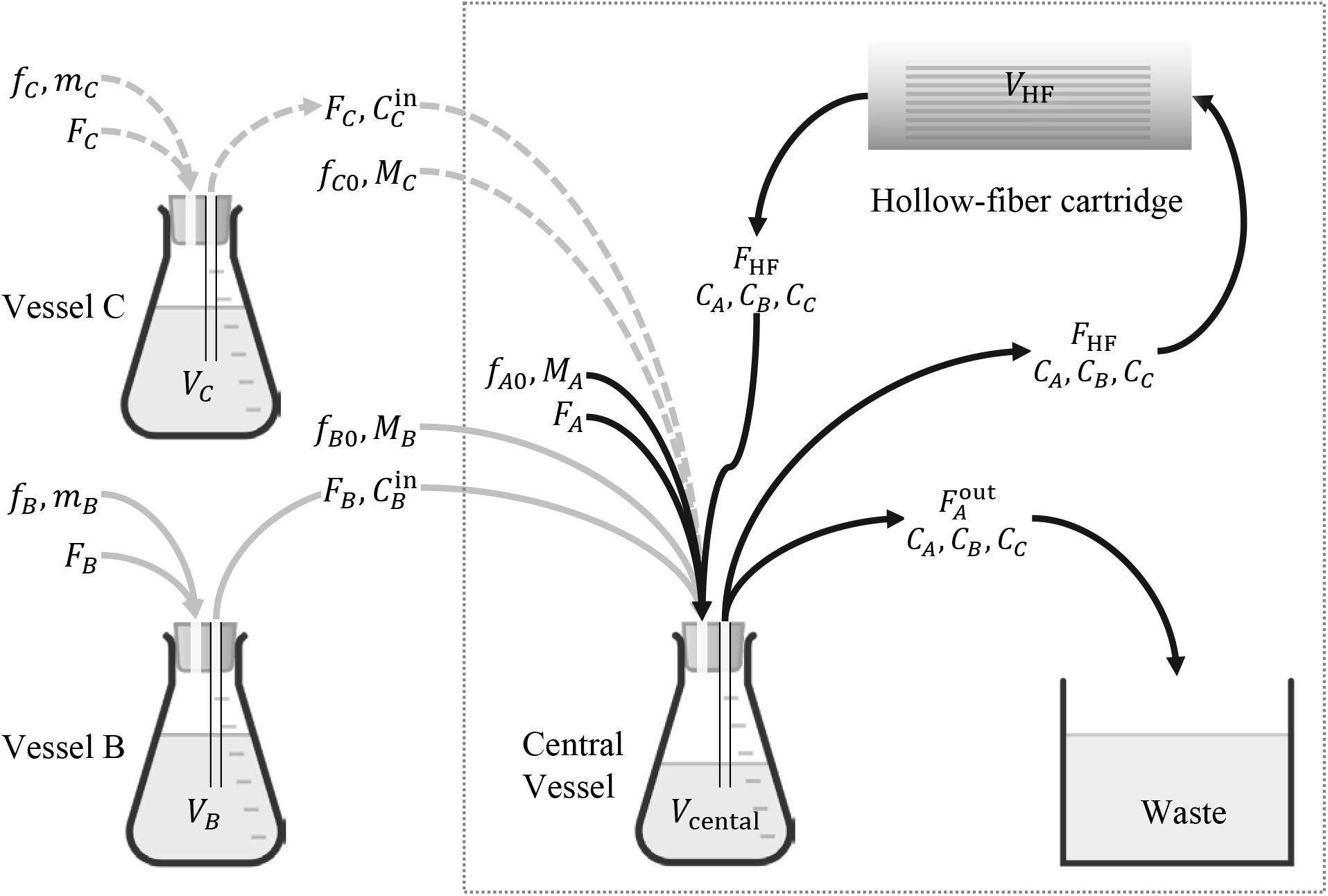
An in-series hollow-fiber system for in vitro simulation of three antimicrobial agents, A, B, C, each following its own elimination pharmacokinetics as it goes through the hollow-fiber cartridge. The notation followed is similar to Fig. 1.

**Fig. 5.**
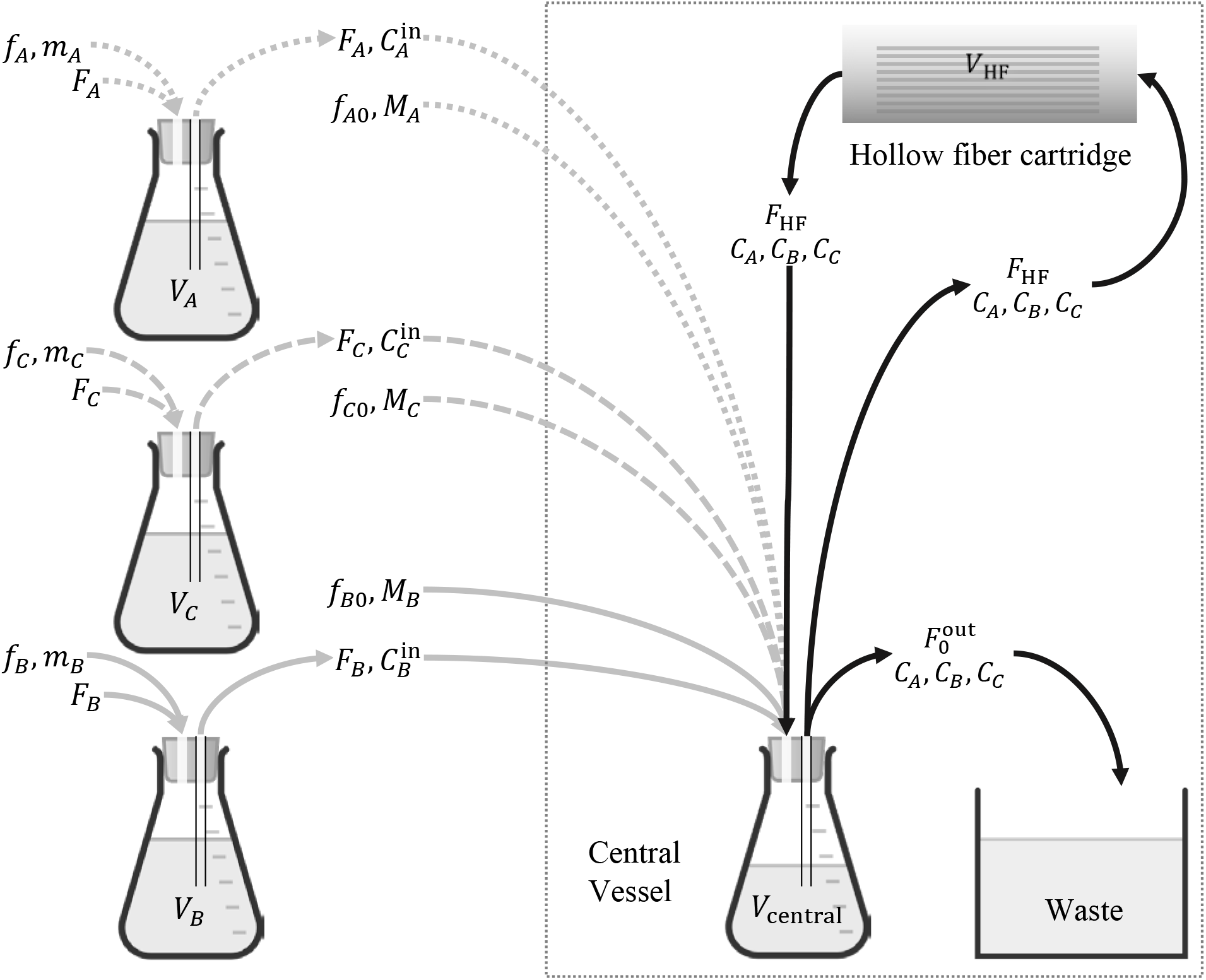
An in-parallel hollow-fiber system for in vitro simulation of three antimicrobial agents, A, B, C, each following its own elimination pharmacokinetics as it goes through the hollow-fiber cartridge.

### Ensuring well-mixed conditions for the central vessel / hollow-fiber cartridge

It is noted that the flow rate *F*_HF_ of the fluid circulating out of the central vessel through the hollow-fiber cartridge and back is assumed to be high enough to ensure that the system comprising the central vessel and hollow-fiber cartridge is well mixed with uniform concentration of each agent throughout. This ensures that all bacteria are exposed to approximately the same concentrations of agents at any time *t*. A quantitative criterion that determines the range of values of *F*_HF_ that are high enough to ensure well-mixed conditions is developed in the Results section.

### Design tasks for in-series and in-parallel configurations

The in-series and in-parallel configurations shown in Fig. 4 and Fig. 5, respectively, require selection of corresponding growth media and agent flow rates as well as liquid volumes in each vessel, such that they can generate exponentially decaying concentrations *C_j_*(*t*) with different half-lives, *t*_1/2*j*_, *j* = *A, B, C*,…, (eqn. (1)). The settings for the corresponding design problems are shown in Table 3 and Table 4. Formulas for completing the system design are subject to analysis. The results of such analysis are presented in the Results section.

**Table 3.**
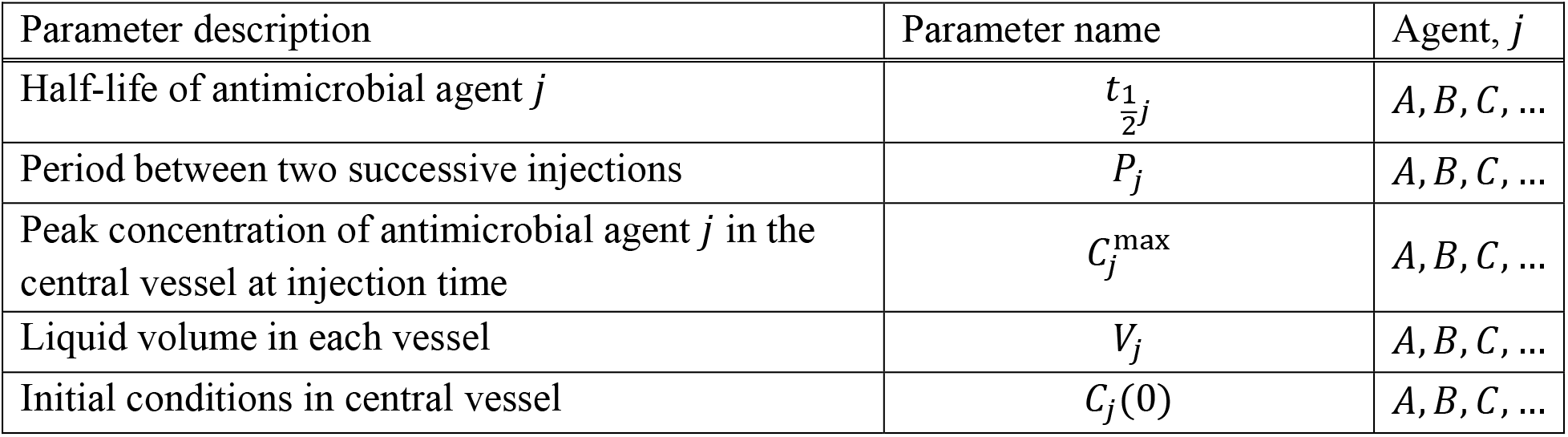
Input data in the design procedure for the in-series hollow-fiber system in Fig. 4 with three (or more) antimicrobial agents

| Parameter description | Parameter name | Agent, $j$ |
| --- | --- | --- |
| Half-life of antimicrobial agent $j$ | $t_{\frac{1}{2}j}$ | $A, B, C, \dots$ |
| Period between two successive injections | $P_j$ | $A, B, C, \dots$ |
| Peak concentration of antimicrobial agent $j$ in the central vessel at injection time | $C_j^{\max}$ | $A, B, C, \dots$ |
| Liquid volume in each vessel | $V_j$ | $A, B, C, \dots$ |
| Initial conditions in central vessel | $C_j(0)$ | $A, B, C, \dots$ |

**Table 4.** Input data in the design procedure for the in-parallel hollow-fiber system in Fig. 5 with three antimicrobial agents

| Parameter description | Parameter name | Agent, $j$ |
| --- | --- | --- |
| Half-life of antimicrobial agent $j$ | $t_{\frac{1}{2}j}$ | $A, B, C, \dots$ |
| Period between two successive injections | $P_j$ | $A, B, C, \dots$ |
| Peak concentration of antimicrobial agent $j$ in the central vessel at injection time | $C_j^{\max}$ | $A, B, C, \dots$ |
| Liquid volume in all vessels | $V_j$ | $A, B, C, \dots$ |
| Liquid volume in combined central vessel and hollow-fiber cartridge | $V_0 \stackrel{\text{def}}{=} V_{\text{central}} + V_{\text{HF}}$ | |
| Initial conditions in central vessel | $C_j(0)$ | $A, B, C, \dots$ |

### Mathematical model for in-series hollow-fiber system

Assuming constant volumes *V_B_, V_C_*,… and negligible flow rate for *f_j_* compared to *F_j_*, mass balance on agent *j, j = B, C*,… around each corresponding vessel in Fig. 4 yields

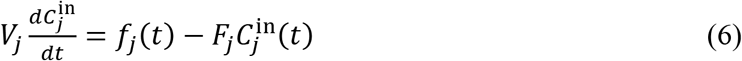

where *j = B, C*,….

Similarly, mass balance on all agents *j = A, B, C*,… around the combined central vessel of effective volume

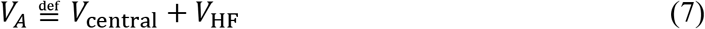

yields

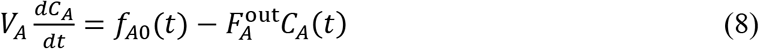

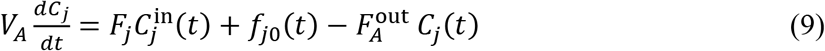

where *j = B, C*,…,

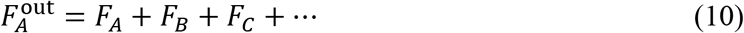

and the effective volume of the central vessel along with the hollow fiber cartridge and the connecting tubes, is assumed to be a well mixed system for high enough circulation rate *F*_HF_, as we show rigorously in APPENDIX A.

The design task entails use of the above mathematical model along with the input data in Table 3 to determine the functions *f*_*j*0_(*t*), *j* = *A, B, C*, …; the functions *f_j_*(*t*), *j* = *A, B, C*, …; and values for the parameters *F_j_, j* = *A, B, C*,….

### Mathematical model for in-parallel hollow fiber system

Assuming constant volumes *V_A_, V_B_, V_c_*,… and negligible flow rate for *f_j_* compared to *F_j_*, mass balance on agent *j, j = A, B, C*,… around each corresponding vessel in Fig. 5 yields eqn. (6) for *j = A, B, C*,… . Similarly, mass balance on all agents *j = A, B, C*,… around the combined central vessel yields

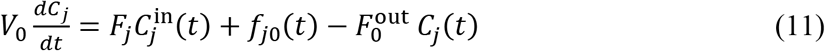

where *j = A, B, C*,…, and

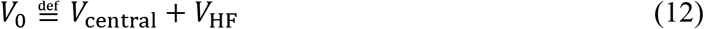

is the effective volume of the central vessel along with the hollow fiber cartridge and the connecting tubes, assumed to be a well mixed system for high enough circulation rate *f*_HF_, as we show rigorously in APPENDIX A.

The design task entails use of the above mathematical model along with the input data in Table 4 to determine the functions *f_j0_*(*t*), *j* = *A, B, C*,…; the functions *f_j_*(*t*), *j* = *A, B, C*,…; and values for the parameters *F_j_*, *j* = *A, B, C*,… .

### Laplace transforms

Development of the proposed experiment design method using the preceding mass balance equations along with the specifications of the design is greatly facilitated by use of Laplace transforms [19] which turn all differential equations of the mathematical problem into algebraic equations that can be easily manipulated to produce the results sought.

### Antimicrobial agents

Levofloxacin powder was acquired from Johnson and Johnson Pharmaceutical Research and Development, LLC. Ceftazidime powder were obtained from Chem-Impex International (Wood Dale, IL). Meropenem powder was purchased from TCI America (Portland, OR).A stock solution in sterile water has been prepared ahead of time and stored in-70°C. Prior to each experimental study, the antimicrobial agents were thawed and diluted to the appropriate concentrations.

### Hollow fiber experiment

The target half-lives and maximum concentrations simulated for meropenem, ceftazidime, and levofloxacin are shown in Table 5. Each drug was dosed at time 0 h. Meropenem and ceftazidime were dosed again at approximately 16 h. Each dose was given 30 minutes to ensure ample mixing in the system. Samples were taken at approximately 1, 2, 3, 4, 6, 8, 17, 18, 19, 20, 22 and 24 h. The liquid volume of the system comprising the central vessel and the hollow-fiber cartridge (eqn. (7)) was *V_A_* = 180 ml with *V*_central_ = 60 ml and *V*_HF_ = 120 ml. The liquid volumes in vessels A and B as well as the injection periods *P_A_, P_B_, P_C_* are also shown in Table 5. The values of *C_A_*(0), *C_B_*(0), *C_C_*(0), were equal to zero. Antimicrobial agent concentrations in the samples were assayed using validated methods by liquid chromatography tandem mass spectroscopy (LC-MS/MS) [20,21].

**Table 5.** Values of pharmacokinetic parameters and vessel volumes for experiment design

| Agent, $j$ | $t_{1/2j}$<br>(h) | $C_j^{\max}$<br>($\frac{\mu\text{g}}{\text{ml}}$) | $V_j$<br>(ml) | $P_j$<br>(h) |
| --- | --- | --- | --- | --- |
| Meropenem (A) | 1 | 10 | 180 | 16 |
| Ceftazidime (B) | 2.5 | 10 | 270 | 16 |
| Levofloxacin (C) | 8 | 10 | 396 | 24 |

Note that the target elimination half-lives mimic human pharmacokinetics. An arbitrary *C*^max^ was used as a proof-of concept. The *C*^max^ observed in humans after clinical dosing can be achieved by a proportional adjustment of the drug amount to inject in the model.

## Results

### Circulation rates for the central vessel / hollow-fiber cartridge system to be well mixed

It is intuitively clear that a high circulation flow rate *F*_HF_ from the central vessel through the hollow-fiber and back (Fig. 4) creates in effect a well mixed system comprising the central vessel and the hollow-fiber cartridge, with effective volume *V_A_*, as shown in eqn. (7). However, it is not immediately obvious how high the flow rate *F*_HF_ should be, to be high enough for reasonable application of the “well mixed” assumption. This question is addressed rigorously in APPENDIX A, where we show that values of *F*_HF_ about 20-30 times the value of 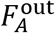 are practically sufficient.

### Design method for in vitro simulation of distinct pharmacokinetics for *N* agents combined: In-series configuration

Given the data in Table 3 and using the mass balances and Laplace transforms as described in Materials and Methods, we show in APPENDIX B that the functions *f*_*j*0_(*t*), *j* = *A, B, C*,… and *f_j_*(*t*), *J* = *B, C*, … can be selected to be impulses of corresponding magnitudes *M_j_* and *m_j_*, i.e.

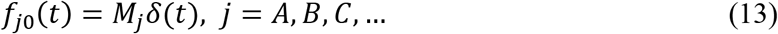

and

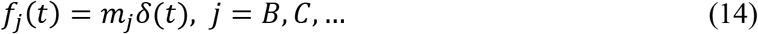

Formulas for values of the parameters *m_j_, j* = *B, C*,…, and *M_j_, j* = *A, B, C*,…, as well as for all flow rates *F_j_*, *j* = *A, B, C*,…, are shown in Table 6. All proofs for the results in Table 6 are shown in APPENDIX B.

**Table 6.** Summary of design procedure given data in Table 3 for the in-series hollow-fiber system in Fig. 4 with three or more antimicrobial agents

| Parameter description | Parameter name and design equation | Agent, $j$ |
| --- | --- | --- |
| Total volumetric flow rate out of central vessel, $F_A^{\text{out}}$ | $F_A^{\text{out}} = V_A k_A$ | |
| Volumetric flow rate, $F_j$, of $j$ into and out of vessel $j$ | $F_j = V_j k_j$ | $B, C, \dots$ |
| Volumetric flow rate, $F_A$, of A into central vessel | $F_A = F_A^{\text{out}} - F_B - F_C - \dots$ | |
| Mass, $M_j$, of bolus injected into the central vessel as pulse over short time $\Delta t$ | $M_j = (C_j^{\text{max}} - C_j(0)) V_A$ | $A, B, C, \dots$ |
| Mass, $m_j$, of bolus injected into the vessel $j$ as pulse over short time $\Delta t$ | $m_j = C_j^{\text{max}} V_A \left( \frac{t_{1/2}^j}{t_{1/2}^A} - 1 \right) = C_j^{\text{max}} V_A \left( \frac{k_A}{k_j} - 1 \right)$ | $B, C, \dots$ |

### Design method for in vitro simulation of distinct pharmacokinetics for *N* agents combined: In-parallel configuration

Similarly, given the data in Table 4 and using the mass balances and Laplace transforms as described in Materials and Methods, we show in APPENDIX C that the functions *f*_*j*0_(*t*), *j* = *A, B, C*,… and *f_j_*(*t*), *j* = *A, B, C*,… can be selected to be impulses of corresponding magnitudes *M_j_* and *m_j_*, i.e.

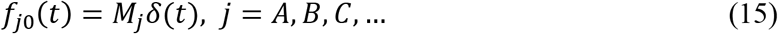

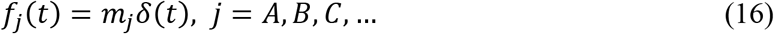

Formulas for values of the parameters *m_j_* and *M_j_, j = A, B, C*,…, as well as for all flow rates *F_j_, j = A, B, C*,…, are shown in Table 7. All proofs for the results in Table 7 are shown in APPENDIX C.

**Table 7.**
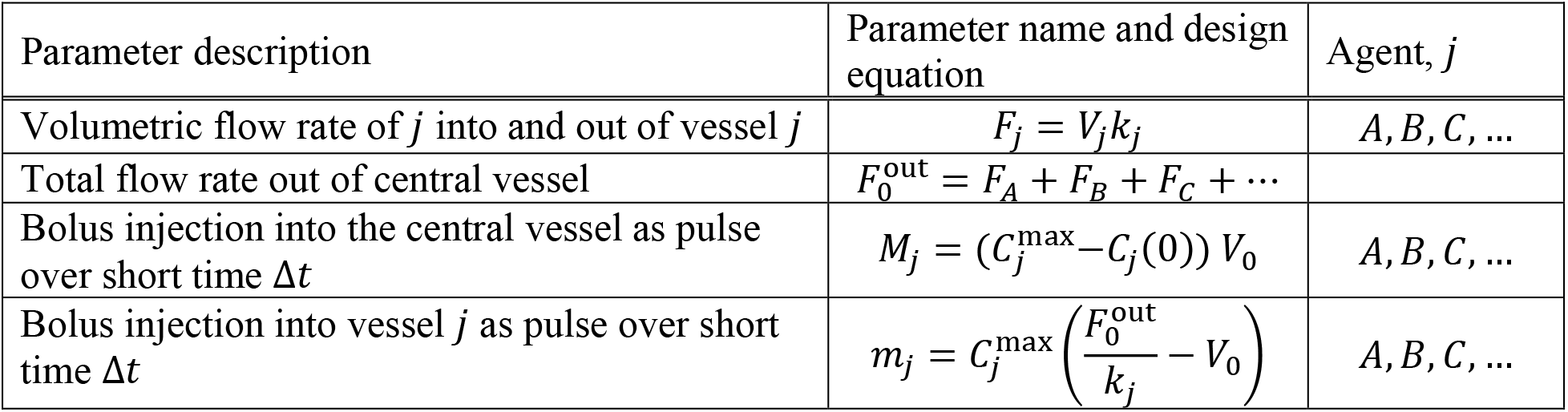
Summary of design procedure given data in Table 4 for the in-parallel hollow-fiber system in Fig. 5 with three or more antimicrobial agents

| Parameter description | Parameter name and design equation | Agent, $j$ |
| --- | --- | --- |
| Volumetric flow rate of $j$ into and out of vessel $j$ | $F_j = V_j k_j$ | $A, B, C, \dots$ |
| Total flow rate out of central vessel | $F_0^{\text{out}} = F_A + F_B + F_C + \dots$ | |
| Bolus injection into the central vessel as pulse over short time $\Delta t$ | $M_j = (C_j^{\text{max}} - C_j(0)) V_0$ | $A, B, C, \dots$ |
| Bolus injection into vessel $j$ as pulse over short time $\Delta t$ | $m_j = C_j^{\text{max}} \left( \frac{F_0^{\text{out}}}{k_j} - V_0 \right)$ | $A, B, C, \dots$ |

### Experiment design resulting from application of developed in-series design method

Using design specifications as shown in Table 5 and with experiment parameters as discussed in the Hollow fiber experiment section, we obtained the following values (Table 8) of the design parameters (Table 6) involved in the in-series experiment configurations (Fig. 4) tested in our laboratory.

**Table 8.** Values of design parameters for in-series configuration of experiment set-up

| Agent, $j$ | $F_j$<br>$\left(\frac{\text{ml}}{\text{min}}\right)$ | $F_A^{\text{out}}$<br>$\left(\frac{\text{ml}}{\text{min}}\right)$ | $M_j$<br>($\mu\text{g}$) | $m_j$<br>($\mu\text{g}$) |
| --- | --- | --- | --- | --- |
| Meropenem, A | 0.26 | 2.08 | 1800 | NA |
| Ceftazidime, B | 1.25 |  | 1800 | 2700 |
| Levofloxacin, C | 0.57 |  | 1800 | 12600 |

### Experiment outcomes

Measurements of agent concentration over time and corresponding model predictions are shown in Fig. 6 for the in-series configuration.

**Fig. 6.**
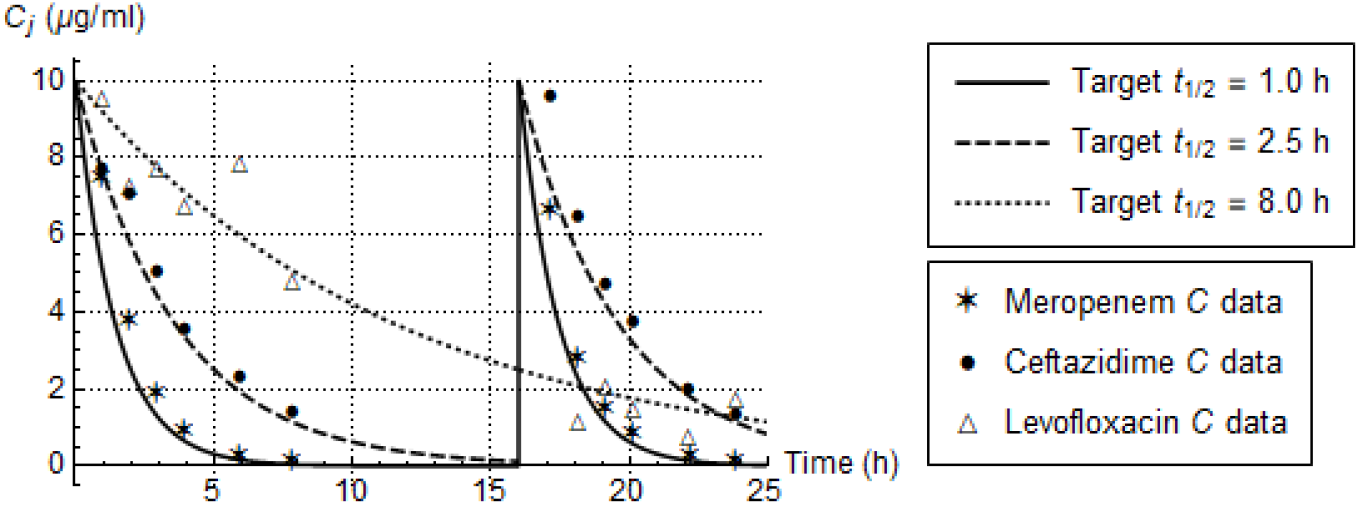
Measurements of concentrations *C_j_*(*t*) for three antimicrobial agents and comparison with model predictions for the in-series experimental set-up shown in Fig. 4.

## Discussion

### Feasibility constraints of in-series design for *N* agents

Given that *m_j_* > 0, *j* = *B, C*,…, in Table 6, the design equation

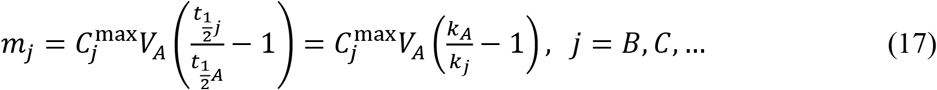

immediately places the constraint

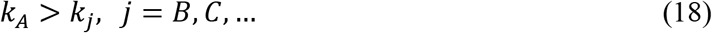

on the selection of agent A, among A, B, C,…, for the central vessel.

In addition, because *F_A_* > 0, the design equation

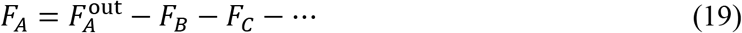

in Table 6 implies

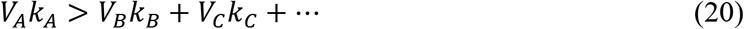

Combined, eqns. (18) and (20) define the feasible area for the design parameters of the in-series configuration.

Note that eqn. (18) is the generalization of eqn. (4), stated by Blaser [12] for 2 agents.

### Comparison of proposed in-series designs to prior design for two agents

When the in-series design for *N* agents, summarized in Table 6, is applied to *N* = 2 agents, it offers additional flexibility compared to the design summarized in Table 2, in a number of ways, as follows:

a. While Blaser’s [12] two-agent design (Table 2) requires that

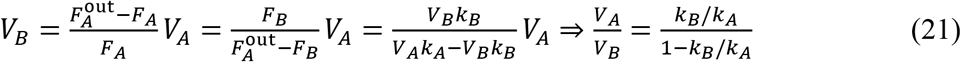

the A-agent design in Table 6 can begin with any given set of volumes *V_j_* (Table 3) as long as the constraints in eqns. (18) and (20) are satisfied. This adds an important degree of freedom to the proposed design. For two agents, A, B, it is interesting to visualize the feasible area of *V_A_, V_B_*, given *k_A_, k_B_*, with *k_A_ > k_B_*, suggested by the *N*-agent design in Table 6, as shown in Fig. 7. That Figure includes Blaser’s [12] two-agent design (Table 2) which is based on eqn. (21).
b. The injection period, *P_j_*, for each antimicrobial agent in all vessels does not have to be large enough to ensure that *C_j_*(*nP*) ≈ 0, *n* = 0, 1, 2,… . Therefore, the values of *C_j_*(0), *j* = *A, B, C*,…, are added to input data shown in Table 3.
c. Unlike Blaser [12], the in-series design in Table 6 does not require that

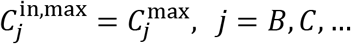

because there is flexibility in the ratios of vessel volumes. Of course, *C_j_*(*t*) and *C*_in_(*t*) share the same exponential decline rate *k_j_*, as explained in APPENDIX B. To visualize the situation for two agents, Fig. 8 shows feasible values of the ratio

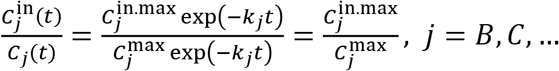

for feasible values of the ratio 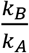 in (0,1). That Figure includes Blaser’s [12] two-agent design (Table 2) which is a single horizontal line at 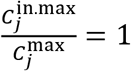. To visualize the situation in yet another way, Fig. 9 shows the profiles of both 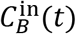 and *C_B_*(*t*) resulting from Blaser’s design [12] in (Table 2) and from the in-series design for *N* = 2 in Table 6 for a few feasible design choices on *V_B_* satisfying 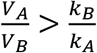 (Fig. 7).

**Fig. 7.**
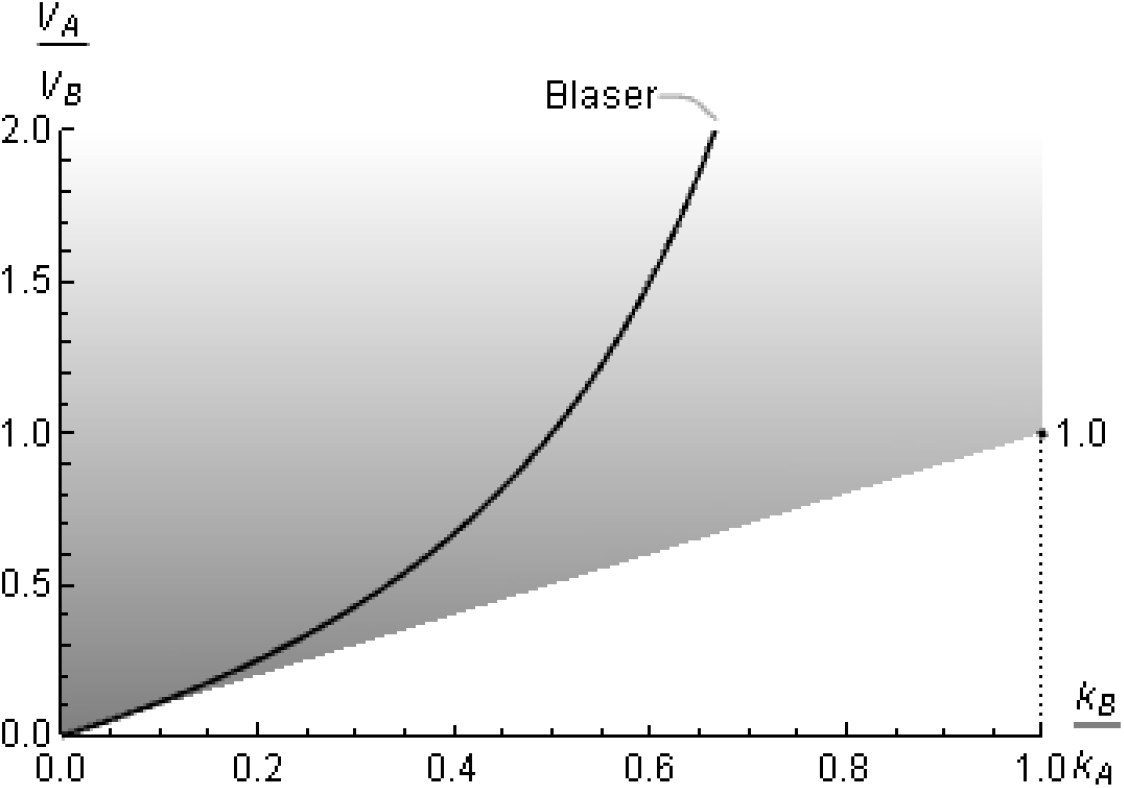
Feasible area (shaded, extending upwards) for 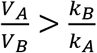 given 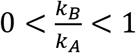 for the in-series design procedure of Table 6 applied to N = 2 agents. Note the design curve according to Blaser [12], following eqn. (21).

**Fig. 8.**
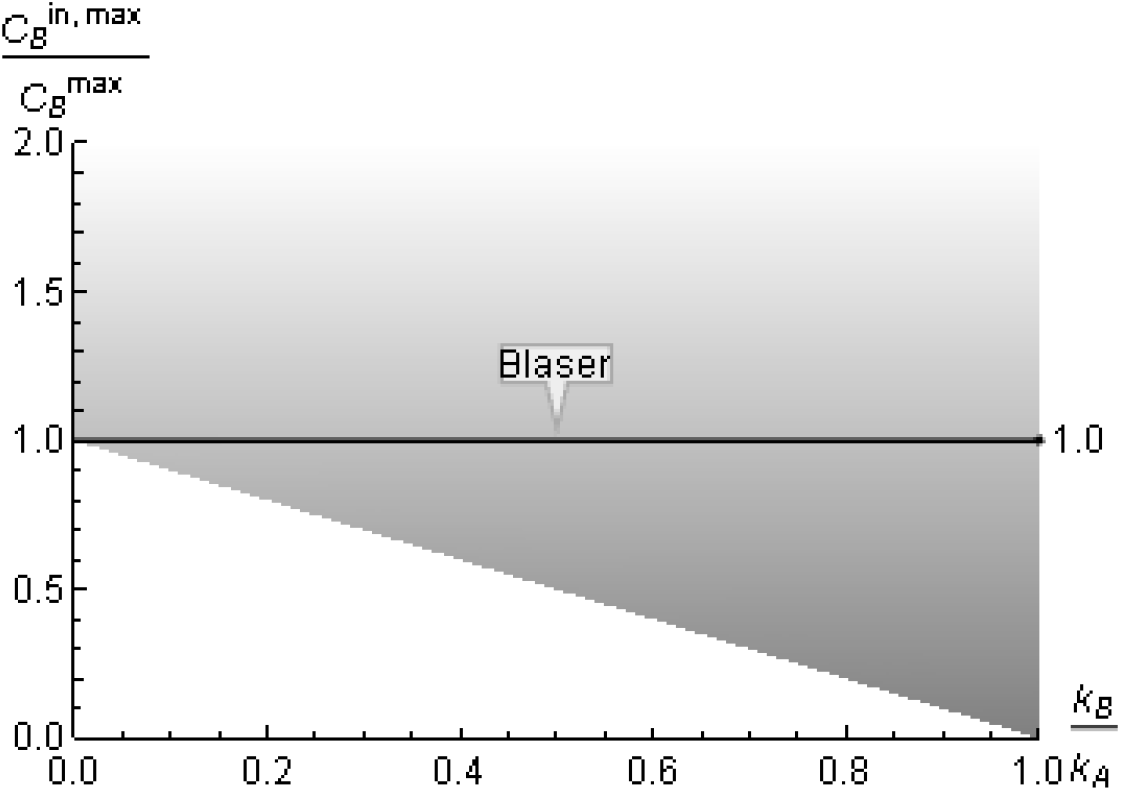
Feasible area (shaded, extending upawrds) for 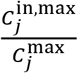 given 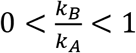 for the in-series design procedure of Table 6 applied to N = 2 agents. Note the fixed value at 1 according to Blaser [12].

**Fig. 9.**
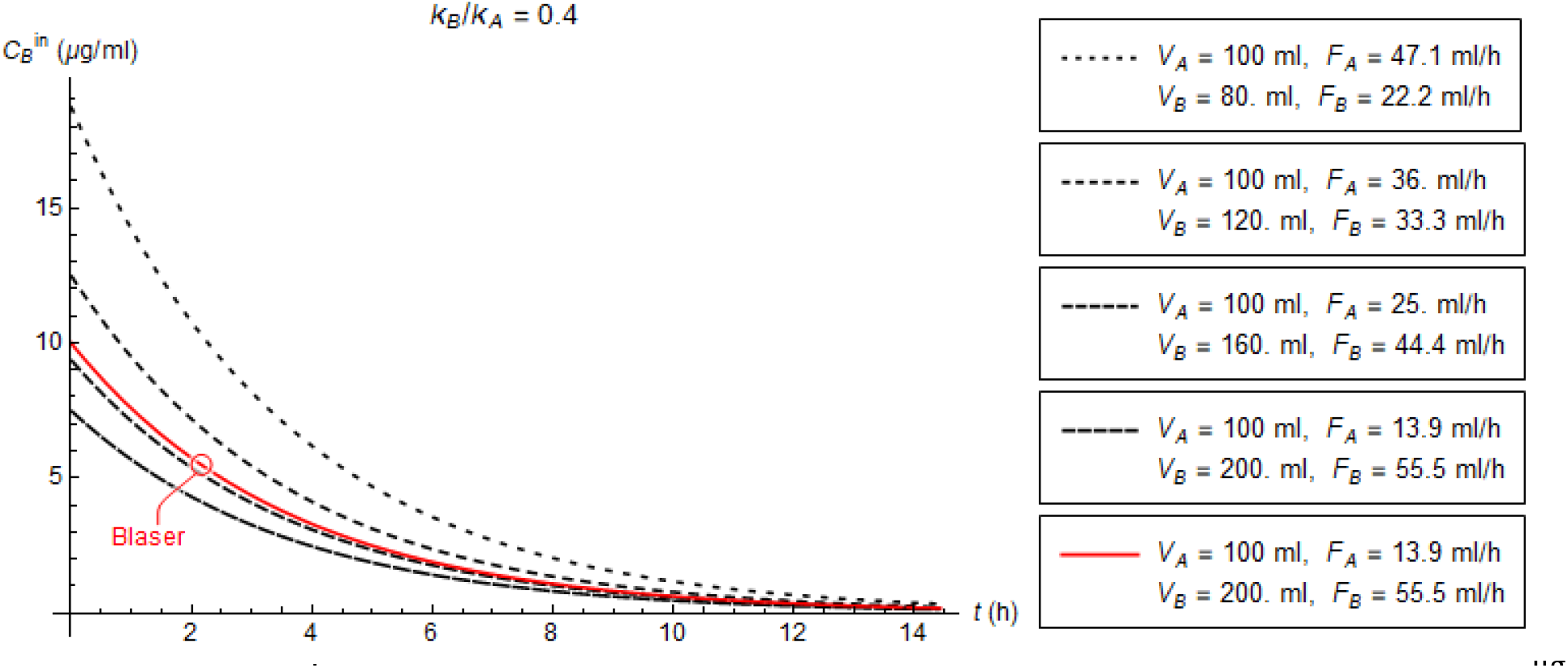
Profile of 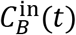 for the system in Fig. 1 with specifications V_A_ = 100 ml, 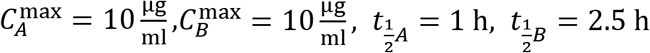, and design parameters {F_A_, F_B_, V_B_, M_A_, M_B_, m_B_} computed according to Blaser [12] (Table 2) and Table 6 for 2 agents. The values of {M_A_, M_B_, m_B_} are {1000,1000,1500} μg for all designs.

We also note that in-parallel designs may result in profile differences between 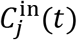 and *C_j_*(*t*) in an entirely similar fashion.

### Feasibility constraints of in-parallel design for *N* agents

Given that *m_j_* >0, *j* = *A, B, C*,…, in Table 7, the design equation

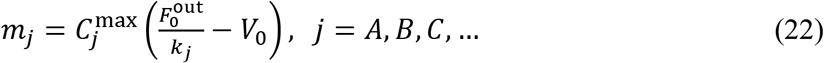

immediately places the constraint 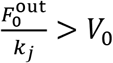, *j* = *A, B, C*,…, which results in the design constraint

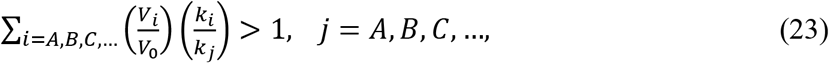

The great importance of eqn. (23) for the flexibility of the in-parallel method is that large enough volumes *V_j_* can always be found, regardless of the values of *k_j_*, *j* = *A, B, C*,…, of *V*_0_, and of any preferable ratios of any one vessel volume over another. How large is large enough can be visualized in the case of two agents, for which eqn. (23) becomes

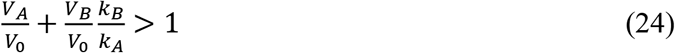

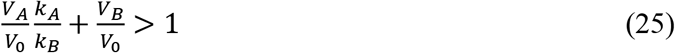

The feasible volume corresponding to eqns. (24) and (25) is shown in Fig. 10.

**Fig. 10.**
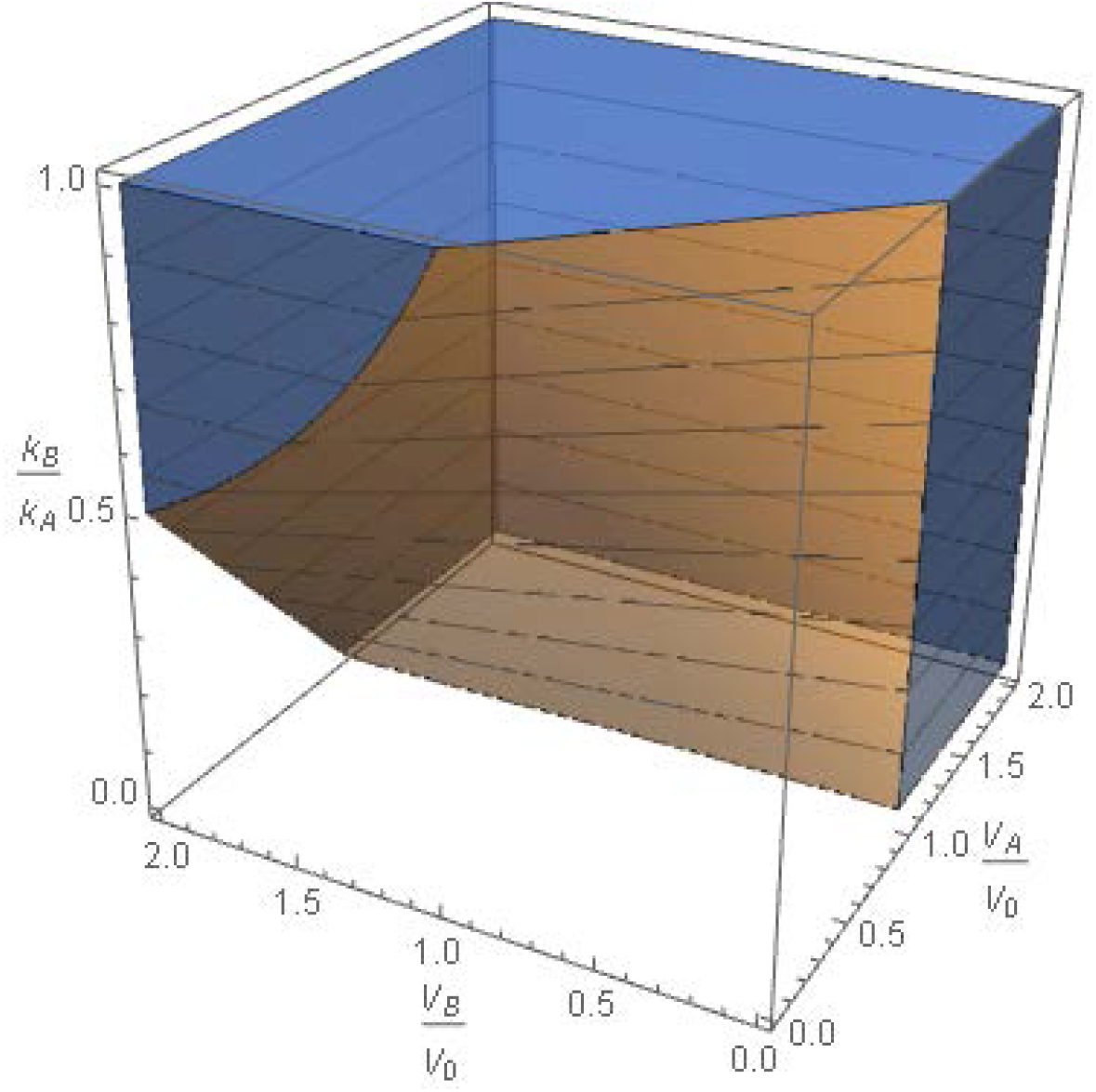
Feasible space for 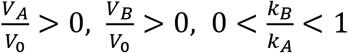 according to eqns. (24) and (25). Note that for 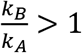 the same plot can be used by swapping A with B, because of symmetry.

## Conclusions

A general method was developed to design an in vitro model for simultaneous simulation of the kinetics of an arbitrary number of *N* drugs with different half-lives. The method developed entails two possible configurations: (a) An in-series configuration, which generalizes for *N* drugs Blaser’s two-drug design [12] and offers additional flexibility even for two drugs, and (b) an in-parallel configuration, which is new and offers different flexibility compared to the in-series configuration. Corresponding design equations were developed for sizing and operation of each configuration (Table 6 and Table 7). The in-series design equations were used for experimental verification using a combination of three antibiotics with distinctly different half-lives (meropenem, ceftazidime, and levofloxacin). Additional experiments in the future would further test the range of applicability of the proposed method. While experimental verification in this work involved antibiotics, the method is applicable to any anti-infective and anti-cancer drugs with distinct elimination pharmacokinetics. With increasing importance of in vitro simulation of the kinetics of an arbitrary number of drugs in combination, the methods developed here are an important new tool for the design of such in vitro models.

## APPENDIX A. High circulation rate implies a well-mixed system

Mass balance for agent A around the central vessel (Fig. 4) immediately yields

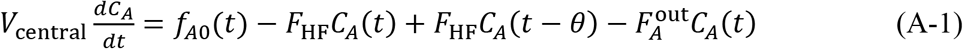

where

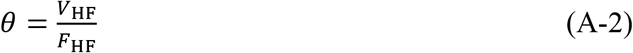

is the time needed for liquid drawn from the central vessel to circulate in plug flow at rate *F*_HF_ through the hollow-fiber cartridge volume *V*_HF_.

Taking Laplace transforms of the above eqn. (A-1) and solving for 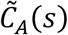 yields

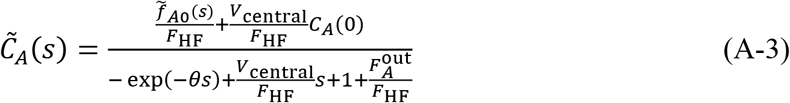

where tilde denotes the Laplace transform of a corresponding function of time. The above eqn. (A-3) implies that *C_A_*(*t*) will include a weighted sum of exponentials of the form *e^Pnt^*, where *p_n_* are the poles of 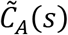, namely the roots of the transcendental algebraic equation

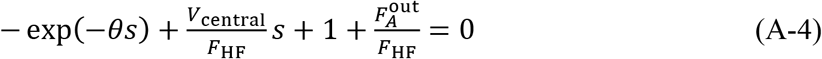

It can be shown that the solution of the above eqn. (A-4) can be expressed in terms of the Lambert function as

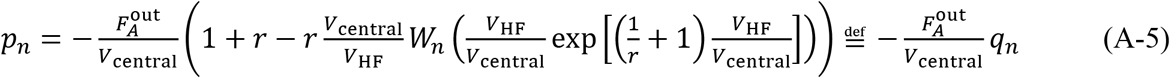

where

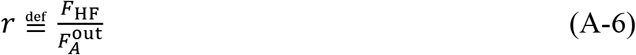

and *W_n_*(*z*) is the Lambert function of order *n* = −1,0,1,2,…, defined as a solution of the algebraic equation *xe^x^* = *z*. Based on the properties of Lambert functions, it is straightforward to show that all *p_n_* have negative real part for *n* = −1,0,1,2,…, therefore *C_A_*(*t*) remains bounded at all times. In addition, all *p_n_* are complex, except *p*_0_. More importantly, Fig. 11 shows that for comparable values of *V*_HF_ and *V*_central_, if *r* is high enough, the real parts of all complex-valued *p_n_* are much larger than the magnitude of *p*_0_, suggesting that the corresponding terms exp(*p_n_t*), *n* ≠ 0 will decay much faster than exp(*p*_0_*t*), so as to be negligible. Therefore, for high enough *r*, the effective volume of the combined system comprising the central vessel and the hollow-fiber cartridge will be

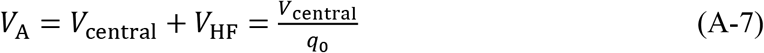

as shown in Fig. 11.

**Fig. 11.**
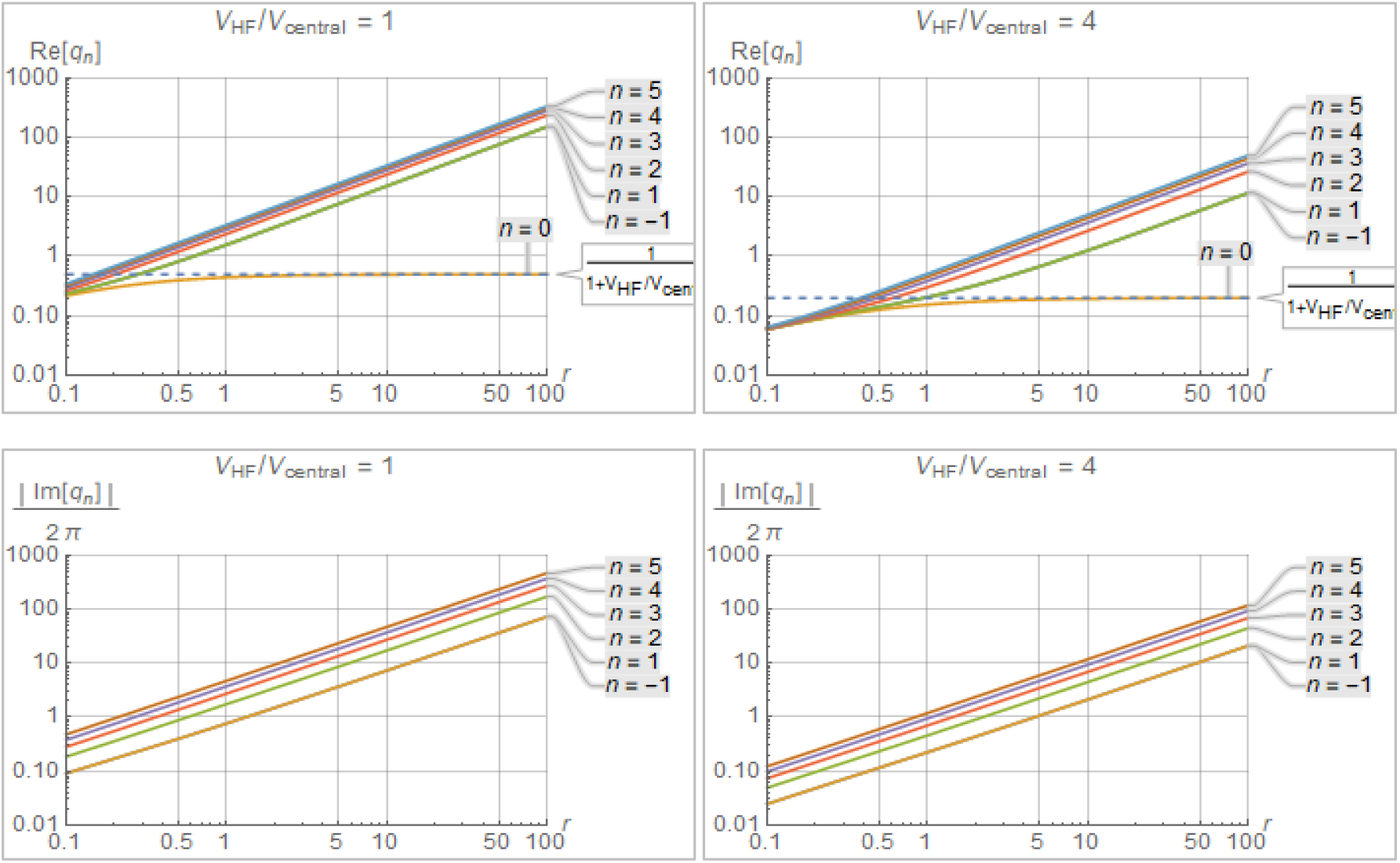
Real and imaginary parts (top and bottom, respectively) of modes 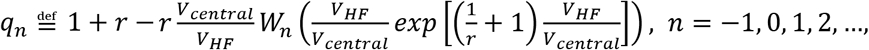 in eqn. (A-5) in terms of circulation ratio 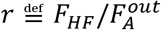. Note that for r greater than about 30, the real part of q_n_, n = −1, 0,1, 2,… suggests that the decay of all 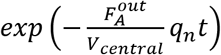, n ≠ 0 is so much faster than the decay of 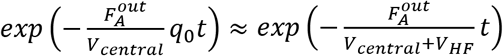 as to be negligible· Furthermore, for r greater than about 30, the imaginary part of q_n_, n = − 1,1, 2,… suggests that no appreciable oscillations are going to appear in 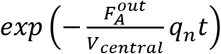, n ≠ 0 as all oscillation frequencies 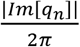, n = −1,1, 2, … are comparable to the exponential decay rates Re[q_n_].

Similar arguments can be made for the agents *B, C*,… going through the hollow-fiber system shown in Fig. 4. Starting with mass balance

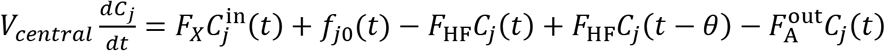

for agent *j* = *B, C*,… one can immediately get

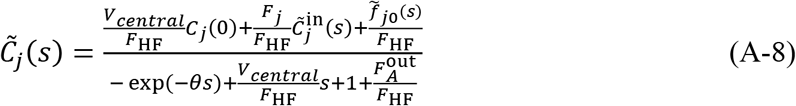

whose denominator is exactly the same as that of 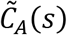 in eqn. (A-3).

The proof for the system shown in Fig. 5 is entirely similar and omitted for brevity.

## APPENDIX B. Proof of eqns. in Table 6

### • Agent A

Taking Laplace transform of eqn. (8) yields

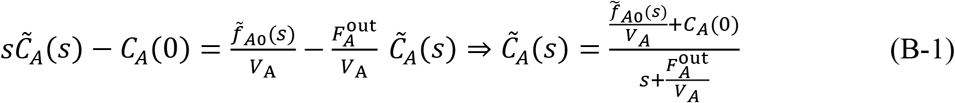

Now, because the concentration of agent A flowing out the central vessel must follow eqn. (1) for *j = A*, it follows that

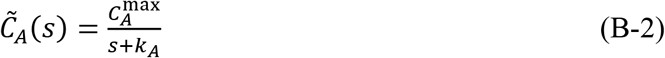

Comparison of the numerators in eqn. (B-1) and (B-2) immediately implies that

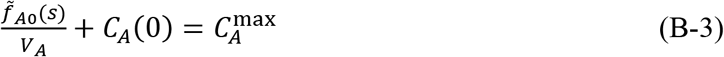

which suggests that 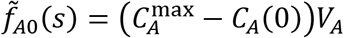, therefore *f*_*A*0_(*t*) must be an impulse of magnitude 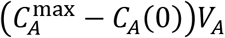:

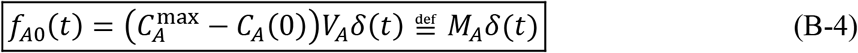

where

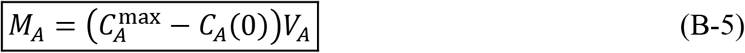

Similarly, comparison of the denominators in eqn. (B-1) and (B-2) immediately implies

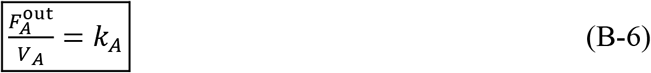

### • Agent *j* = *B, C*,…

Following the same approach as above for agent *j, j = B, C*,…, one can take Laplace transform of eqn. (9) to get

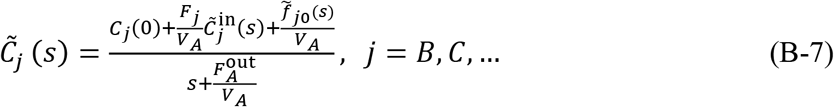

Because the concentration of agent *j* flowing out the central vessel must follow eqn. (1) for *j* = *B, C*,…, it follows that

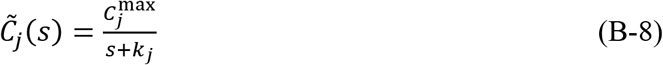

Comparison of eqns. (B-7) and (B-8) immediately implies that the term (*s* + *k_j_*) must be introduced into and the term 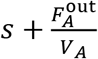 eliminated from the denominator of the right side of eqn. (B-7). To accomplish this, a simple and experimentally practical choice is to select

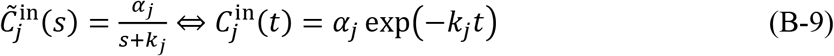

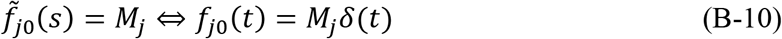

and to require that

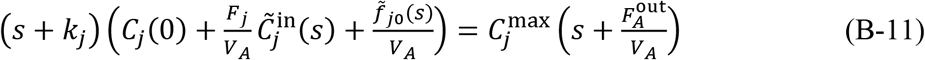

The exponential decline in eqn. (B-9) implies that the appropriate profile of 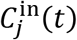 over time can be easily constructed by setting

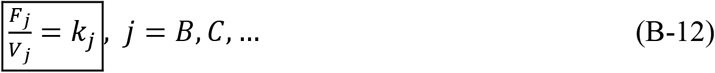

The values of the parameters *α_j_* and *M_j_*, needed to complete the above design of 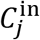 and *f*_*j*0_, can be easily determined by matching like powers of s in eqn. (B-11). This, combined with eqn. (B-10) yields

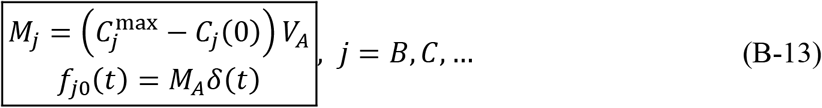

and combined with eqn. (B-9) yields

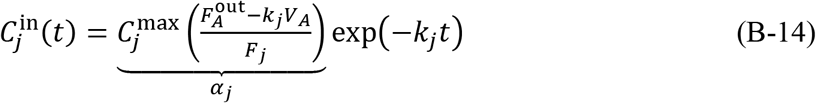

The above profile for 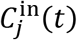 can be easily created by the corresponding to an injection of bolus

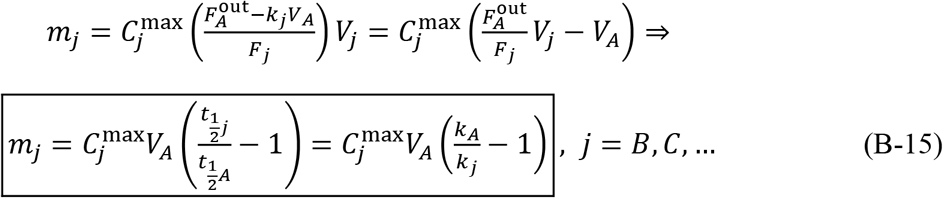

into vessel *j = B, C*,… .

## APPENDIX C. Proof of eqns. in Table 7

Taking Laplace transform of eqn. (9) for the symbols used in Fig. 5 yields

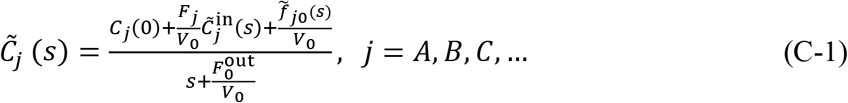

Now, because the concentration of agent *j* flowing out the central vessel must follow eqn. (1) for *j = A, B, C*,…, it follows that

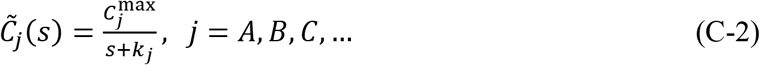

Comparison of eqns.(C-1) and(C-2) immediately implies that

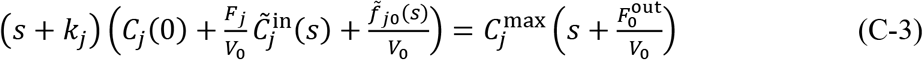

In entirely similar fashion as in APPENDIX B, one gets

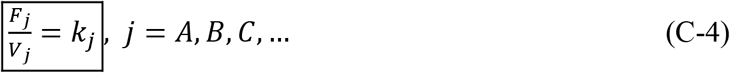

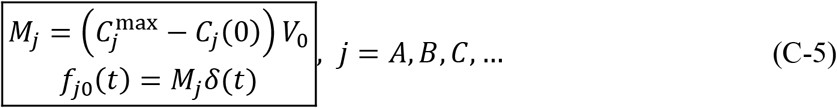

and

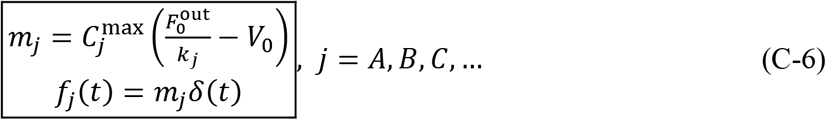

